# Depth relationships and measures of tissue thickness in dorsal midbrain

**DOI:** 10.1101/2020.05.13.093492

**Authors:** Paulina Truong, Jung Hwan Kim, Ricky Savjani, Kevin R. Sitek, Gisela E. Hagberg, Klaus Scheffler, David Ress

**Affiliations:** Department of Neuroscience, Baylor College of Medicine, Houston, TX, 77030 USA; Department of Neuroscience, Rice University, Houston, TX, 77005 USA; Department of Radiation Oncology, University of California, Los Angeles, CA, 90095 USA; High Field Magnetic Resonance, Max Planck Institute for Biological Cybernetics, Tübingen, Germany; Department of Biomedical Magnetic Resonance, Eberhard Karl’s University of Tübingen and University Hospital, Tübingen, Germany

**Author notes:** Corresponding Author: 1 Baylor Plaza T115E, Houston, TX 77030, USA.

**Keywords:** MRI, depth, midbrain, colliculus, periaqueductal gray

## Abstract

Dorsal human midbrain contains two nuclei with clear laminar organization, the superior and inferior colliculi. These nuclei extend in depth between the superficial dorsal surface of midbrain and a deep midbrain nucleus, the periaqueductal gray matter (PAG). The PAG, in turn, surrounds the cerebral aqueduct (CA). This study examined the use of two depth metrics to characterize depth and thickness relationships within dorsal midbrain using the superficial surface of midbrain and CA as references. The first utilized nearest-neighbor Euclidean distance from one reference surface, while the second used a level-set approach that combines signed distance from both reference surfaces. Both depth methods provided similar functional depth profiles generated by saccadic eye movements in a functional MRI task, confirming their efficacy for superficial functional activity. Next, the boundaries of the PAG were estimated using Euclidean distance together with elliptical fitting, indicating that the PAG can be readily characterized by a smooth surface surrounding PAG. Finally, we used the level-set approach to measure tissue depth between the superficial surface and the PAG, thus characterizing the variable thickness of the colliculi. Overall, this study demonstrates depth-mapping schemes for human midbrain that enables accurate segmentation of the PAG and consistent depth and thickness estimates of the superior and inferior colliculi.

## 1 INTRODUCTION

Human midbrain mediates a panoply of critical brain functions that range from homeostasis to perception to cognition. However, studying the human dorsal midbrain, including the superior colliculus (SC) and inferior colliculus (IC), is challenging due to the small size and deep location of its component nuclei, as well as limited methods for understanding their internal structure. As a result, most of our existing knowledge about the function of the midbrain arises from research conducted in animal models. As human ultra-high field MRI adoption grows, the constraints posed by collicular size and location are reduced due to increased MR signal at 7T and above. However, the challenge of understanding the internal structure of the midbrain remains, requiring a depth-mapping scheme for probing the anatomy and function of the human midbrain.

Both SC and IC have an intricate laminar cytoarchitecture and exist in a complex 3D topology that is folded into four small hillocks on the dorsal surface of the midbrain. The inner boundaries of the colliculi abut another nucleus, the periaqueductal gray (PAG), which surrounds the ventricular cerebral aqueduct (CA).

The laminar functional organization of mammalian colliculi has been clearly established based on several electrophysiology and lesion studies in animal models. Experiments in non-human primates and cats have shown that the layers of SC can be divided into three groups with independent functional purposes. Superficial layers have been shown to receive direct visual input and have retinotopically organized receptive fields (Cynader & Berman, 1972; Feldon & Kruger, 1970). Intermediate layers mediate oculomotor control (Robinson, 1972), while deep layers are associated with multimodal inputs, primarily multisensory and visuomotor neurons (Meredith & Stein, 1986; Sprague & Meikle, 1965). Likewise, electrophysiological studies in IC of rats and primates have shown that the central nucleus organizes auditory inputs of varying frequency in a laminar fashion (Baumann et al., 2010; Schreiner & Langner, 1997). Invasive studies have exhibited this tonotopic organization along the dorsal-to-ventral direction, which roughly corresponds to laminar depth (Baumann et al., 2011; Cheung et al., 2012; Loftus, Malmierca, Bishop, & Oliver, 2008; Malmierca et al., 2008; Schreiner & Langner, 1997)

While functional MRI (fMRI) has been used extensively in human cerebral cortex, the human brainstem has been relatively neglected despite its critical role in brain function. Functional contrast-to-noise ratio (CNR) is often low due to its deep location (Singh, Pfeuffer, Zhao, & Ress, 2017) and functional data is susceptible to partial volume effects due to the small size of important brainstem structures compared to conventional fMRI voxel sizes of 3–4 mm. However, several recent high-resolution fMRI studies using millimeter-scale voxels have demonstrated detailed analysis of the functional laminar topography of the human colliculi with reasonable CNR. Indeed, these studies have confirmed the functional properties of the three major layer groups of SC (Katyal & Ress, 2014; Linzenbold & Himmelbach, 2012; Loureiro et al., 2017; Savjani, Katyal, Halfen, Kim, & Ress, 2018; Schneider & Kastner, 2010), as well as the auditory organization of IC (De Martino et al., 2013; Moerel, De Martino, Uğurbil, Yacoub, & Formisano, 2015; Ress & Chandrasekaran, 2013), encouraging interest in resolving these laminar variations of human colliculi in more detailed experiments. A recent fMRI study showed depth-dependent variations of the BOLD response with 1-mm sampling in SC (Loureiro et al., 2017). This work utilized binary mask erosion techniques to generate depth-dependent regions-of-interest (ROI). The approach resolves depth at the 1-mm scale but does not permit associations between laminae at different depths.

In previous work, we utilized a nearest-neighbor Euclidean point-to-surface definition of depth to define layers in SC by creating depth kernels that associate a particular locus of superficial tissue with deeper tissue. These kernels were small computational cylinders extending from the surface of the brainstem inwards, normal to the surface of SC. With this method, we successfully demonstrated depth-dependent BOLD responses from visual stimulation and attention (Katyal & Ress, 2014; Katyal, Zughni, Greene, & Ress, 2010) as well as saccadic activity in SC (Savjani et al., 2018) and frequency-dependent auditory stimulation in IC (Ress & Chandrasekaran, 2013).

However, defining a laminar neighborhood of tissue using nearest-neighbor associations is a drastic oversimplification. In the small convoluted colliculi, the cylinders are susceptible to oversampling or undersampling deep tissue, depending on the gradient of the curvature. In order to capture the physical structure of collicular layers as observed in histology and MR microscopy, a more topologically consistent method of defining depth within the colliculi is desirable. Several methods have been developed to compute depth in cortex, including solutions of Laplace’s equation (Jones, Buchbinder, & Aharon, 2000) and equi-volume methods (Waehnert et al., 2014). However, they are computationally expensive to solve in a stable and accurate fashion and have never been applied to midbrain.

Here, we compare two methods for depth-analysis in human midbrain: our previous Euclidean approach, and an algebraic level-set algorithm that utilizes two surfaces. We adapted the level-set method originally implemented in cortex (Kim, Taylor, & Ress, 2017) to human midbrain using the superficial brainstem tissue-CSF boundary as the outer surface and the CA as the inner surface to create a depth coordinate normalized to the thickness of the tissue under investigation. First, we compared the ability of both methods to delineate visual- and saccade-evoked function in superficial SC, concluding that both methods perform similarly. We also delineated the inner boundaries of the colliculi in order to set a precedent for future functional studies in the deepest layers of SC. This was accomplished by localizing the outer boundary of the PAG using a combination of Euclidean and level-set methods applied to gray-matter tissue-probability maps obtained at 9.4T. Lastly, we evaluated anatomical thickness of the colliculi, using both methods to measure the distance from the collicular surface to the outer boundary of the PAG. In this case, the level-set approach provided more precise measures of tissue depth in the superior and inferior colliculus. Altogether, we show that our level-set method for depth analysis meets or exceeds the performance of more typical Euclidean approaches and suggests new applications of our method for mapping the human brainstem.

## 2 METHODS

### 2.1 – Subjects

We applied our level-set scheme to twenty subjects. Eight of these participated in subcortical fMRI experiments at 3T (Savjani et al., 2018), giving informed consent under procedures reviewed and authorized by the Baylor College of Medicine Institutional Review Board. The other twelve subjects participated in quantitative MRI experiments at 9.4T (Hagberg et al., 2017) and underwent a physical and psychological check-up by a local physician and provided written informed consent following local research ethics policies and procedures. These investigations were conducted in agreement with the World Medical Association Declaration of Helsinki in its most recent version (2013).

### 2.2 – Acquisition & segmentation of anatomical images

For each of the eight subjects scanned at 3T, we acquired high-resolution (0.7-mm voxels) T1-weighted anatomical volumes using an MP-RAGE sequence (TI = 900 ms, TR = 2600 ms, 9° flip angle, TA = 22 min) on a 3T Siemens Trio scanner with a product 32-channel head coil (Erlangen, Germany). The remaining twelve subjects were scanned at 9.4T (Siemens Medical Solutions, Erlangen, Germany) using a 16-channel, dual-row transmit array operating in the circularly polarized mode, and a 31-channel receive array (Shajan et al., 2014). We acquired B_1_ field maps by the actual flip angle imaging method (Yarnykh, 2010; Yarnykh & Yuan, 2004) with nominal flip angle FA=60°; repetition time TR_1_/TR_2_=20/100ms; echo time TE=7 ms, voxel size=3×3×5 mm^3^; and acquisition time TA = 225 s. We further acquired high-resolution (0.8-mm isotropic voxels) quantitative whole brain T_1_-maps using an MP2-RAGE sequence (TI1/TI2 = 900/3500 ms, TR = 6 ms, volume TR = 9 s, TE = 2.3 ms, GRAPPA = 3, partial-Fourier factor 6/8, 256 RF pulses, and TA = 9 min 40 s). At 9.4T, this approach permitted high-quality anatomical imaging with the much shorter acquisition time than at 3T. The MP2RAGE images were reconstructed off-line while correcting for deviations from the nominal excitation flip angle and for T_2_-dependent deviations in inversion efficiency of the adiabatic inversion pulse (Hagberg et al., 2017). From the quantitative T_1_ maps, two sets of synthetic, B_1_-artifact-free, T_1_-weighted, MP2RAGE contrast images were generated pixel-wise from the analytical model equations described in the Appendix of Marques et al. (2010). The first image set was used for brain tissue segmentation in volume space using SPM12. The white-matter grey matter tissue contrast could be increased with respect to the image acquisition by setting the TI2 to 2650ms, and the inversion efficiency to 0.9 while the remaining parameters were the same as in the MR-acquisition. The second image set was generated to improve the quality of the FreeSurfer segmentation (performed using 100 iterations and the following recon_all options: ‘–highres −3T – schwartzya3t-atlas’) and was based on the following parameters: TI_1_/TI_2_ = 950/2100ms; flip angle = 5/3°; volume TR = 6 s; and an inversion efficiency of 0.8451 (corresponding to the experimentally determined median value found in brain tissue).

For all subjects, to generate the outer segmentation, the brainstem tissue (BS) of each subject was initially identified using a probabilistic Bayesian approach in the FreeSurfer 6.0 software package (Iglesias et al., 2015). These segmentations were then edited manually to obtain precise delineation of the four colliculi and their vicinity. To create the inner segmentation, the CA was identified using a mixture of manual and automatic region-growing tools implemented in ITK-SNAP (Yushkevich et al., 2006).

### 2.3 – Surface modeling

A surface model *S*_1_ was estimated at the interface between the brainstem tissue and adjacent CSF or thalamic tissues (Fig. 1A). An initial isosurface was created directly from the segmentation using MATLAB R2016a (Mathworks Corp., Natick, MA, USA). Then, to reduce voxelation artifacts, we applied five iterations of refinement using a variational, volume-preserving deformable surface algorithm (Bajaj, Xu, & Zhang, 2008; Khan et al., 2011; Xu, Pan, & Bajaj, 2006). The same procedure was applied to the CA segmentation to construct a second surface, *S*_2_. (Fig. 1E).

**Figure 1.**
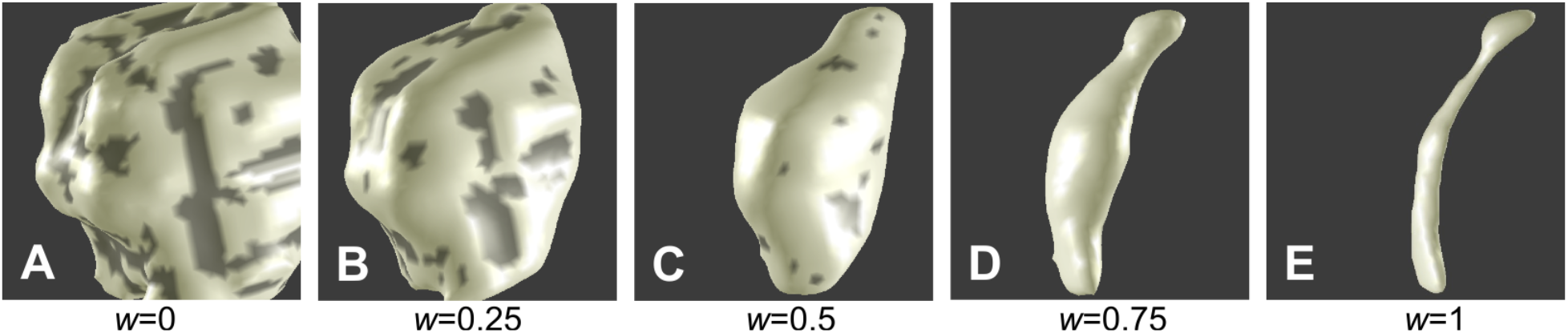
Isosurfaces of tissue proximal to the cerebral aqueduct constructed at five values of *w* for one subject scanned at 3T, with *w*=0 representing the interface between brainstem tissue and CSF (*S*_1_) and *w*=1 representing the cerebral aqueduct (*S*_2_).

### 2.4 – 3D level-set depth-mapping

We calculated a normalized signed distance function *w* using two separate physical distances relative to the corresponding surfaces (Fig. 2A, B). First, Euclidean distances were calculated for voxels in the volume to the nearest triangle of the designated surface *S*_1_ and *S*_2_, giving a 3-dimensional mapping of each voxel to its distance value (Eberly, 1999). Second, the sign for each voxel was determined based on its location; voxels enclosed within the surface were assigned positive distance values, while voxels outside the surface were given negative distances. Thus, distances from *S*_1_ (*d*_1_) became increasingly positive from the surface to BS into deeper tissue, while distances from *S*_2_ (*d*_2_) were only positive within CA and negative in BS and surrounding CSF. Finally, we calculated a weighted sum of the signed distance functions, with distance parameter *w*, using a level-set scheme that solves an Eikonal equation (Bajaj et al., 2008; Khan et al., 2011; Kim et al., 2017):

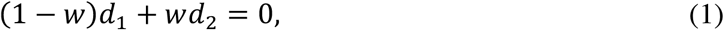

where the level-set parameter *w* provides a normalized depth coordinate between the two surfaces. The depth metric *w* (Fig. 2C) is zero on the brainstem surface (***d*_1_ = 0**) and unity at the surface of CA (***d*_2_ = 0**); *w*< 0 is outside BS and *w*> 1 within CA. Because the normalized depth coordinate is independent of the variable tissue thickness, the above equation also enables generation of isosurfaces that evolve smoothly from *S*_1_ to *S*_2_ (Fig. 1A—E) as well as outside of the domain bounded by the surfaces. Taking advantage of this computational feature, the changes in curvature that distinguish the colliculi became less obvious with increasing *w* (moving from brainstem surface to CA), and the level-set surfaces of *w* ultimately morph into the cerebral aqueduct, *S*_2_. Normalized depth maps were consistent between subjects (Fig. 3), showing a smooth progression between the brainstem surface at *w* = 0 to the CA at *w* = 1. As evident in the surfaces shown in Fig. 1, the characteristic curvature of the colliculi diminished with increasing *w* depth.

**Figure 2.**
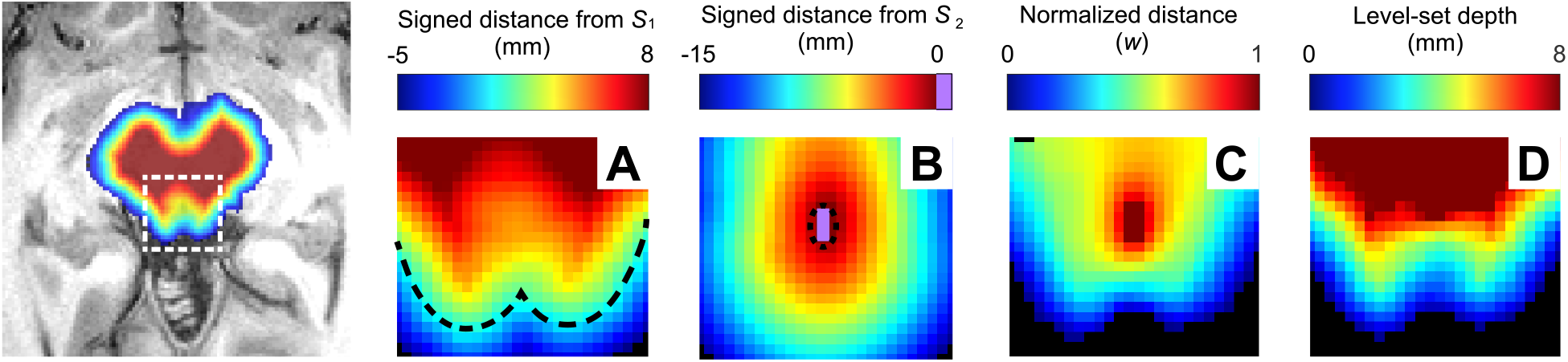
Distance maps generated for one subject overlaid on an axial T1 MRI image of superior colliculus (far left) acquired at 3T. Shown in panels to the right are Euclidean signed distances **(A)** *d*_1_, **(B)** *d*_2_, **(C)** normalized depth, *w* and **(D)** level-set depth derived from *w* by ray tracing. Black dashed lines show the surface representations of the superior colliculus (*S*_1_) in A and the cerebral aqueduct (*S*_2_) in B.

**Figure 3.**
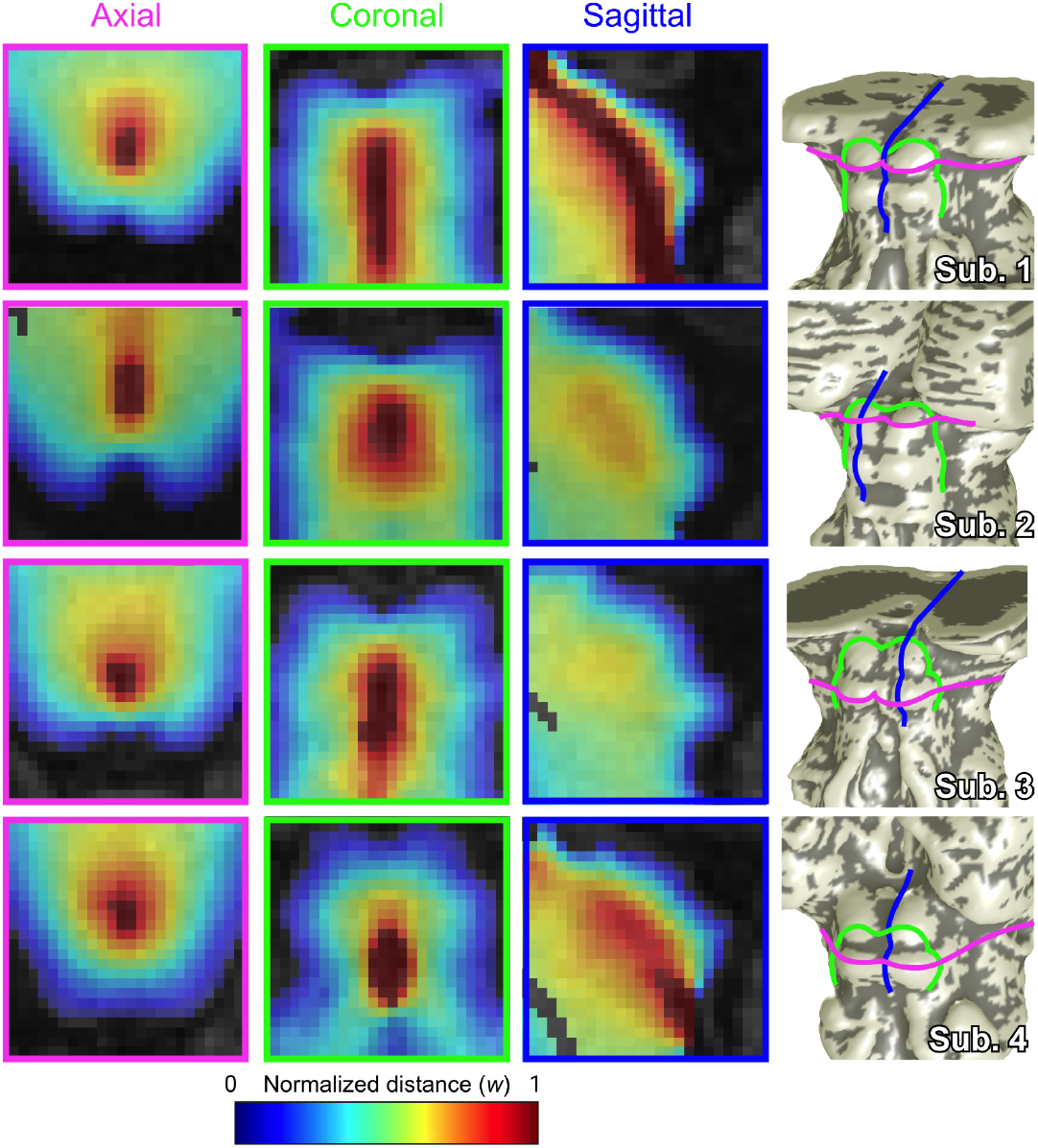
Left to right: axial, coronal, and sagittal cross-sections of normalized depth maps in four subjects scanned at 3T, and location of cross-sections (pink: axial, green: coronal, and blue: sagittal) on brainstem surface models (*S*_1_) of each subject. Axial cross-sections show both superior colliculus (subjects 1 and 2) and inferior colliculus (subjects 3 and 4). Sagittal cross-sections ran through the cerebral aqueduct (subject 1), through the crown of the colliculi (subjects 2 and 3), and through tissue adjacent to the cerebral aqueduct (subject 4).

### 2.5 – Generation of streamlines and depth-averaging kernels

To obtain unique correspondences between *S*_1_ and *S*_2_, we treated *w* as a pseudopotential. We calculated 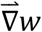 by convolution with 5-point stencil kernels along each dimension. Then, we used ray tracing of 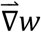 to generate streamlines originating at each vertex of *S*_1_ (Fig. 4A) and propagating throughout midbrain and vicinity. The streamlines were initialized along the corresponding surface normals of *S*_1_ and propagated in small piecewise increments (0.25 voxels), following 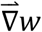 from *w* = 0–1.5. The same process was repeated following the negative gradient from *w* = 0 to *w* = −1. The positive and negative streamline segments were then concatenated into a continuous streamline from *w* = −1 to *w* = 1.5. To avoid numerical pathologies, we utilized several heuristics. Topological inconsistencies were avoided by terminating streamlines that encountered large changes (>80°) in the direction of 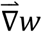 in a single step. Additionally, stagnating streamlines that failed to make sufficient spatial progress after each iteration beyond a minimum threshold (typically 0.05 voxels) before reaching CA were removed. Any streamlines that remained incomplete (*w* < 1) were also discarded after a maximum number of iterations (typically 64 in the direction of positive *w* and 32 in the negative direction).

**Figure 4.**
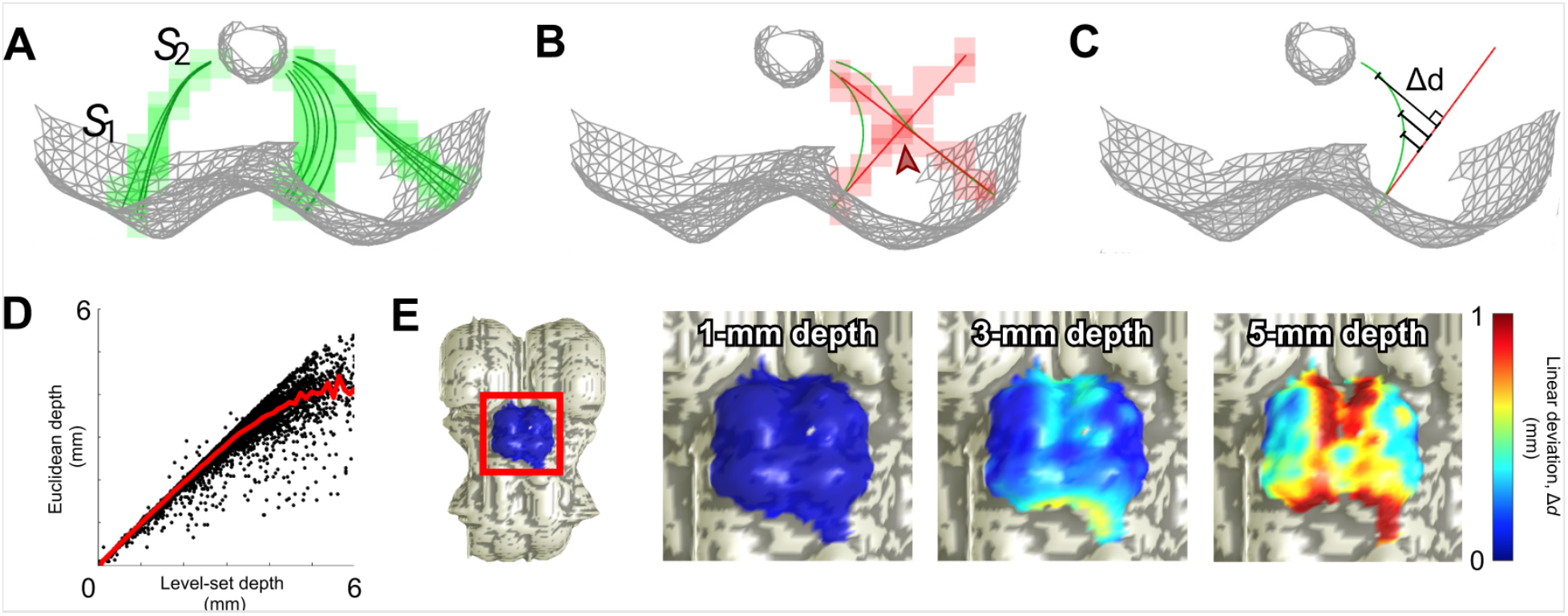
**(A)** Quasi-axial view of a surface representation of superior colliculus (*S*_1_) and cerebral aqueduct (*S*_2_) in one subject scanned at 3T with three level-set depth-averaging kernels (radius=0.7 mm, manifold distance). Their component streamlines (dark green) are regridded (light green) to the spatial resolution of the anatomical volume (0.7 mm isotropic voxels). **(B)** In comparison, cylindrical depth-averaging kernels demonstrate undesirable overlap (“contention”, red arrow) and inappropriately directed sampling in deeper collicular tissue. **(C)** Quantification of the deviation metric, Δ*d*, between level-set and Euclidean sampling methods. **(D)** Relationship between Euclidean nearest-neighbor depth and level-set depth in collicular tissue. **(E)** Surface representations of brainstem (*S*_1_) of one subject, with deviation Δ*d* values at three level-set depths mapped back to the surface vertices from which each streamline originates.

To create depth-averaging kernels that associated voxels on the surface of BS with deeper tissue, we gathered all streamlines within a chosen radius, typically 0.7-mm, of manifold distance of each vertex on *S*_1_ (Fig. 4A). The coordinates of each streamline were then rounded to the nearest voxel and associated with their corresponding vertex of origin on the brainstem tissue surface.

For comparison, we utilized our earlier approach that used only the Euclidean distance metric *d*_1_ to define depth. In this method, depth kernels were computed by extending a 0.7-mm manifold radius disk of BS surface voxels along their mean normal in both directions, yielding a cylinder of voxels (Fig. 4B).

### 2.6 – Depth profiles of visual stimulation-saccade-evoked activity in SC

To test the new level-set streamline approach, we re-analyzed existing data on the polar-angle representation of saccadic eye movements in SC (Savjani et al., 2018). For both stimulation and saccades, we measured the peak BOLD response at depths between 0.5 mm outside BS to 3.5 mm inside BS at increments of 0.1 mm, averaging the values within each 1.2-mm-wide bin. We expected the profiles for saccades to be shifted deeper into SC compared to those from visual stimulation. The reliability of our data was established using a bootstrapping method. We resampled across runs with replacement over 2000 iterations, calculating a new laminar profile for each resampled average. This data was used to obtain the mean depth profiles across subjects with their corresponding 68% confidence intervals, and depth values of peak activity with their 68% confidence intervals and *p*-values (Fig. 5). Depth profiles generated using streamlines and level-set physical tissue depth were compared to profiles using our previous cylinders and Euclidean depth.

**Figure 5.**
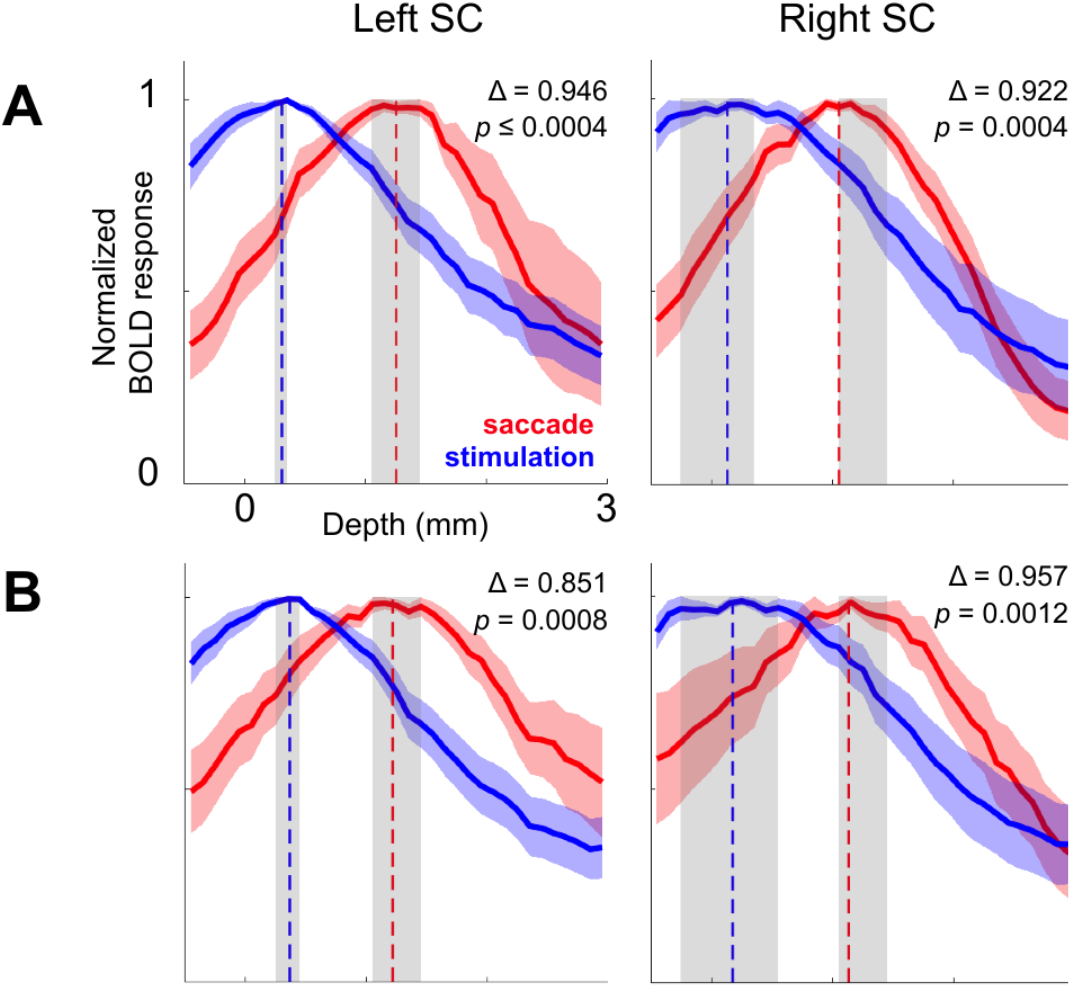
Bootstrapped (*n*=2000) mean laminar profiles of normalized BOLD response in superior colliculus (SC) across 4 subjects scanned at 3T, depth-averaged using Euclidean depth and cylinder depth kernels (**A**) vs. level-set depth and streamline depth kernels (**B**). 68% confidence intervals are shown.

### 2.7 – PAG analysis in midbrain

We estimated the outer boundary of the PAG using quantitative structural data obtained at 9.4T. For this purpose, the unified segmentation approach (Ashburner & Friston, 2005) and a gray-matter tissue-probability atlas (Lorio et al., 2016) in SPM12 were applied to the MP2RAGE-based image sets with optimized white-grey matter contrast (as described above). This map was qualitatively discriminative of the PAG boundary at very high probability thresholds (Fig 6A). To quantify an optimal probability threshold (*P_thr_*), we minimized spatial uncertainty (D. B. Ress, Harlow, Marshall, & McMahan, 2004) of the PAG boundary across subjects. First, we combined our Euclidean distance metric from CA, *d*_2_, with our new level-set depth kernels to quantify *P* as a function of depth, binning increasing depth from *S*_1_ to *S*_2_, at increments of 0.05 mm (Fig. 6B). Though we initially calculated depth profiles using the cylinder depth kernels for comparison, they provided almost no contrast between the PAG and the surrounding tissue. From the streamline depth profiles, we then computed the spatial uncertainty *U*:

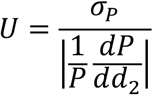

where *σ_P_* is the standard deviation (SD) of *P* across subjects at each value of *d*_2_. Examining the mean values of *U* with respect to gray matter probability, we determined that the optimal *P_thr_*= 0.96. We then constructed and characterized a 3D estimate of the PAG boundary (Fig. 6C, D). The segmentation for CA was upsampled by a factor of 2 and skeletonized to create an axis. Using this axis as a normal vector, 2D slices were taken through the brainstem probability map at each point along the axis. These contours were then fit in a least-squared error sense with ellipses centered around the cerebral aqueduct. These ellipses were characterized by three parameters: left-right axis length *a*, dorsal-ventral axis length *b*, and aspect ratio *ε* = *b/a*. (Fig 6E). These 2D ellipses were then filled in and transformed back to the locations of the slices they were calculated from, to create a 3D segmentation of the elliptical fit of the PAG. We termed the resulting surface generated from this segmentation *S*_ellipse_.

**Figure 6.**
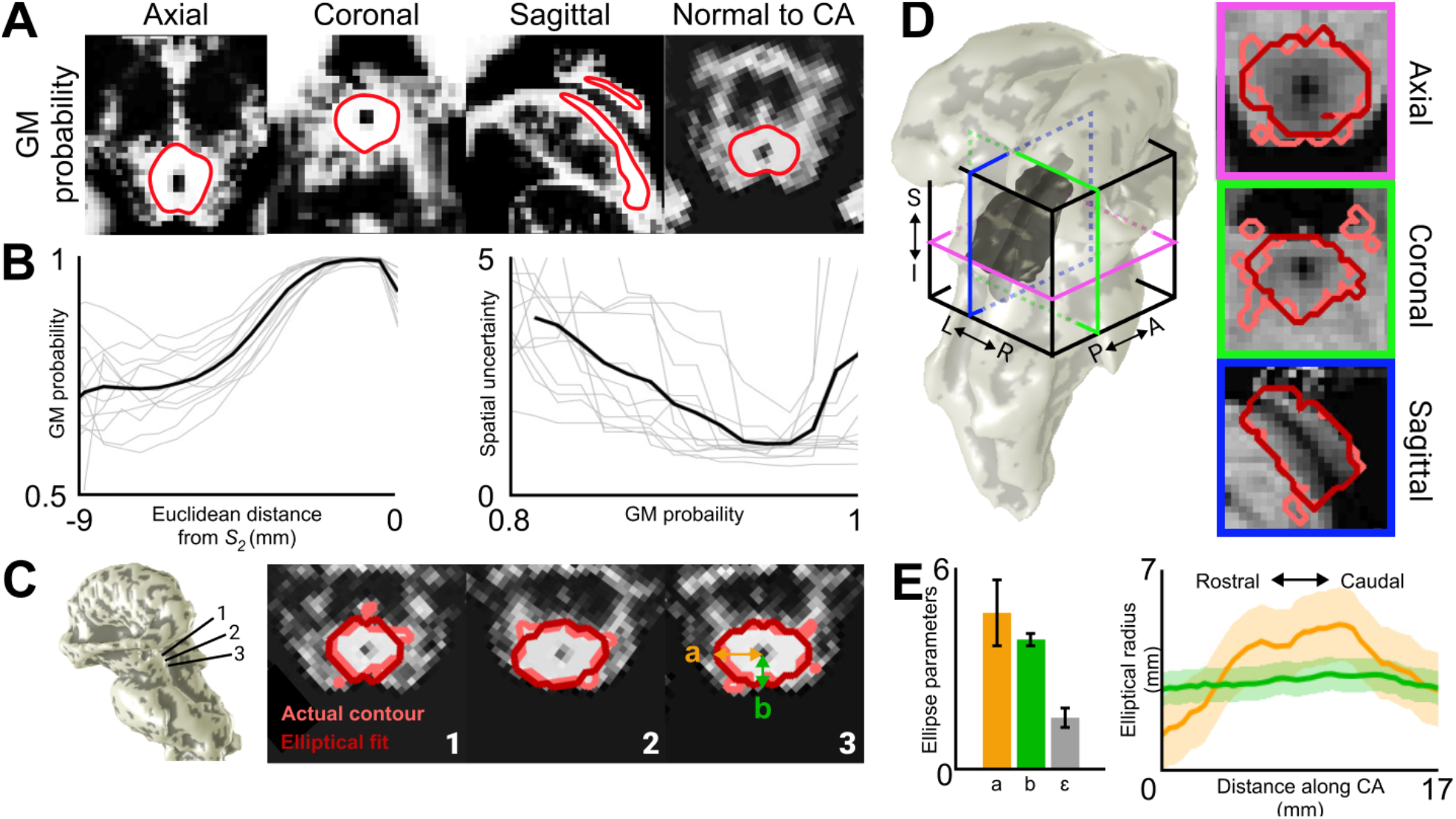
**(A)** Gray-matter probability map of one subject scanned at 9.4T with contour lines drawn at *P*=0.96 (red). **(B)** Left: Individual depth profiles (gray) sampling gray-matter probability maps using streamlines, with respect to distance from cerebral aqueduct (CA). Mean depth profiles across subjects is shown in black. Right: Individual values of spatial uncertainty *U* with respect to gray-matter probability. Mean across subjects is shown in black. **(C)** Slices of the gray-matter probability map of one subject taken normal to CA. Three sections are shown running through superior colliculus (SC; left), inter-collicular sulcus between SC and inferior colliculus (IC; center), and IC (right). Contour lines drawn at *P*=0.96 are shown in light red, and the resulting elliptical fits to these 2D slices are in dark red. **(D)** Surface representations of the PAG (dark gray) enclosing *S*_2_ (CA; light gray), shown with *S*_1_ (brainstem surface; light gray. Three orthogonal slices of one subject’s T_1_ anatomy volumes are shown. Light red depicts the actual PAG contour at *P*=0.96 (*S*_PAG_). Dark red depicts the boundary of the estimated PAG segmentation (*S*_ellipse_) reconstructed from transforming the 2D elliptical fits back to their 3D locations. **(E)** Left: Mean+/-std of elliptical parameters (*a*=left-right semi-axis; *b*=dorsal-ventral semi-axis; *ε*=aspect ratio) across subjects. Right: Mean±SD elliptical parameters *a* (orange) and *b* (green) along the length of the cerebral aqueduct, normalized to the mean length of CA.

A second 3D representation of the PAG was also obtained as a smooth iso-probability surface, *S*_PAG_, of the gray-matter probability map at *P_thr_* using the methods described above (Fig 6D). Upon grossly comparing *S*_PAG_ and *S*_ellipse_, the former was, as expected, very similar to the elliptical fit described above.

### 2.7 – Measurements of collicular thickness

We used path distances along the streamlines to create a new distance map estimating physical tissue depth, obtained by contour integration along each streamline. The path integrals, which oversample the volume, are then re-gridded using a Delaunay triangulation approach (Amidror, 2002) onto the anatomical reference volume (Fig. 2D). This level-set depth stopped increasing at CA, whereas Euclidean depth (Fig. 2A) continued to increase into the center of BS.

For the twelve subjects from which we collected gray-matter probability maps to delineate the PAG, we measured collicular tissue thickness using three different depth metrics: Euclidean distance, level-set path distance, and normalized w distance. Then, collicular thickness was calculated by integrating path distance along each streamline from the surface of the colliculi to *S*_PAG_ (7A). Table 1 displays mean collicular thickness values at six particular locations: the four peaks of left and right SC and IC, and the two sulci between the left and right colliculi of each SC/IC pair. Our method for automatically selecting these points is as follows (Fig. 7B).

**Table 1.** Mean±SD for tissue thickness values in left (L) and right (R) colliculi and inter-collicular sulci (ICS) for SC and IC in all 12 subjects using three different depth metrics.

| Units | SC |  |  | IC |  |  |
| --- | --- | --- | --- | --- | --- | --- |
|  | L | R | ICS | L | R | ICS |
| Physical (Euclidean) | 2.64±1.59 | 3.09±1.83 | 1.04±0.70 | 3.35±0.48 | 3.11±1.17 | 0.88±0.35 |
| Physical (Streamline) | 4.19±2.84 | 4.80±2.79 | 1.23±0.66 | 3.18±0.32 | 2.75±1.04 | 0.93±0.43 |
| Normalized (*w*) | 0.43±0.16 | 0.42±0.13 | 0.26±0.13 | 0.39±0.04 | 0.32±0.12 | 0.20±0.07 |

**Table 2.** Coefficient of variation (COV) of tissue thickness values using three different depth metrics across all 12 subjects.

| Units | SC |  |  | IC |  |  |
| --- | --- | --- | --- | --- | --- | --- |
|  | L | R | ICS | L | R | ICS |
| Physical (Euclidean) | 0.602 | 0.593 | 0.671 | 0.144 | 0.377 | 0.397 |
| Physical (Streamline) | 0.676 | 0.582 | 0.539 | 0.101 | 0.381 | 0.465 |
| Normalized ($w$) | 0.364 | 0.298 | 0.513 | 0.094 | 0.369 | 0.342 |

**Figure 7.**
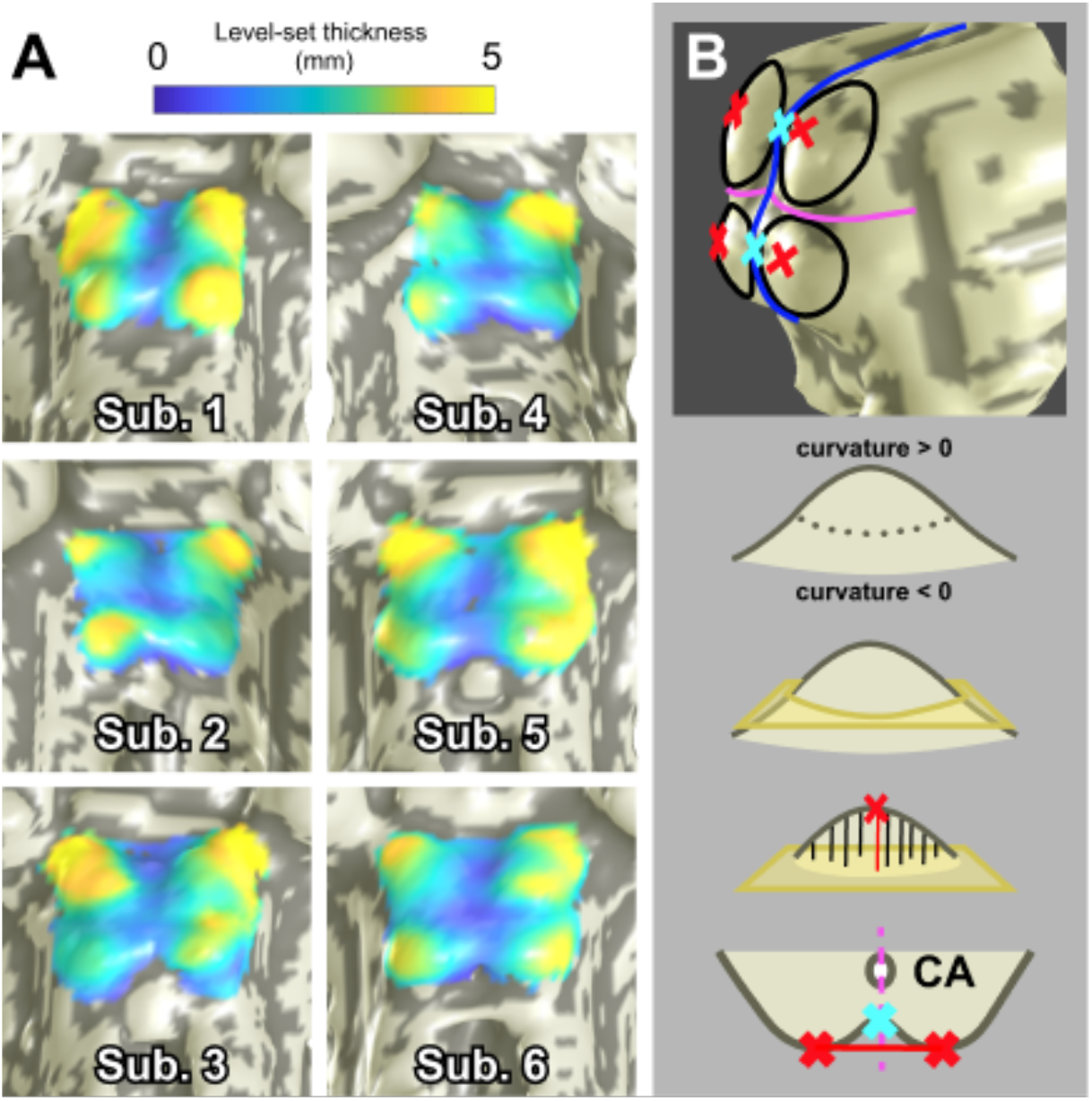
**(A)** Surface representations of brainstem (*S*_1_) of four subjects scanned at 9.4T, overlaid with collicular tissue thickness maps. **(B)** Determination of collicular peaks and inter-collicular sulci. The collicular surface *S*_1*A*_ was divided into four quadrants using a quasi-sagittal (blue) and axial (fuchsia) plane. A base plane (yellow) through the vertices with zero curvature was defined for each colliculus, and collicular peaks (red) were defined to be the vertices with greatest distance to each base plane Inter-collicular sulci (cyan) were vertices on the mid-sagittal plane closest to the line between left and right peaks.

First, for each subject, we isolated the vertices of *S*_1_ of corresponding to the colliculi. We manually created a third segmentation label for the voxels of SC and IC in ITK-SNAP that grossly segregated the surfaces of all four colliculi. We then mapped these voxels by nearest-neighbor association to the BS surface. For completeness, we then expanded this surface by an additional 2-mm manifold distance, thus creating *S*_1*A*_.

Second, *S*_1*A*_ was used to locate the peaks of each colliculus by dividing its vertices into four quadrants, each containing one colliculus, based on a set of planes. A quasi-sagittal plane separating left and right colliculi was specified as the plane of best fit through the vertices of the CA surface model, *S*_2_. To calculate an axial plane separating SC and IC, the collicular surface vertices were divided into bins of 0.5-mm width according to their axial position. The midpoint of the bin with the most negative mean curvature defined the axial plane of separation. We used these planes to divide *S*_1*A*_ into four subsets, each containing a single collicular surface. Next, we defined a base-plane for each colliculus by taking all vertices with positive curvature values and fit these with a plane using a least-squares method. We then defined the peak location as the point with maximum Euclidean distance from this base-plane. We also experimented with defining the collicular peaks as the vertices with greatest positive curvature but found that these points of maximum curvature varied significantly across individual colliculi as well as across subjects. In contrast, the peak distance definition was far more stable, and was therefore used as our standard approach.

Third, for the inter-collicular sulcus (ICS) of SC, we calculated the line between the peaks of left and right SC calculated as described above. The intersection point between this line with the plane-of-best-fit through the CA was located, and the vertex of *S*_1_ closest to this point, within 0.25 mm of the plane of best fit, was defined to be the center of the inter-collicular sulcus. For the inferior sulcus, we repeated the same process with left and right peaks of IC.

Once we obtained the six points of interest, we measured each of their thickness values using three different metrics. For Euclidean distance, we used the distance along the surface normal from *S*_1_ to *S*_PAG_. For level-set physical distance, we used integrated path distance along the streamline originating from each point to the point where it intersected the PAG. For normalized distance, we linearly interpolated the *w*-distance map at the point where the streamlines of interest intersected the PAG. Lastly, we applied a Gaussian-weighted smoothing kernel (2-mm diameter, 1-mm full-width-at-half-maximum [FWHM]) to each of these points to obtain estimates of the six thickness values. For each of *n* vertices within 1 mm of the peak or sulcus in question, its thickness value *t* was given a weight *A* according to its manifold distance *d_m_* to the center point of the kernel: 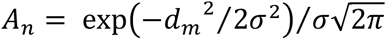, where *σ* is the standard deviation of the Gaussian function, and 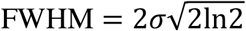. The final thickness value was then calculated as the weighted sum, 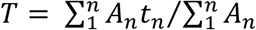.

## 3 RESULTS

### 3.1 – Euclidean vs. level-set methods

Unlike the kernels generated using the cylinder method, the level-set streamlines avoided overlap through their nonlinearity and compression with increased depth (Fig. 4A and B). The streamlines in the colliculi exhibited various degrees of non-linear deformation depending on starting location and their depth into BS. With respect to superficial location, the greatest nonlinearities were expected to occur in streamlines that originate from regions where the curvature varies rapidly (Fig. 4A and B). Low amounts of nonlinearity at the lateral edges of collicular tissue were observed, gradually increasing inwards towards the concave tissue between the colliculi.

With respect to metrics of physical tissue depth, the relationship between Euclidean and level-set depth was sub-linear (Fig. 4D), reflecting increasing curvature in deeper collicular tissue. Additionally, level-set depth, or path distance along the streamlines, stopped increasing at CA, whereas Euclidean depth continued to increase into the center of BS (Figs. 2A and D).

We established a metric Δ*d* to quantify the deviation between level-set and cylinder depths (Fig. 4C). In addition to measuring deviation between the two methods, this comparison also quantified the non-linearity of the level-set depth trajectories. At each point along each streamline, Δ*d* was the orthogonal distance to the *S*_1_ normal vector of the vertex from which the streamline originated. Greater values of Δ*d* at increasing level-set depth were consistent with the warping observed above. Δ*d* was mapped back to *S*_1_ at various depths (Fig. 4E). Non-linearity was very small (≪1 mm) for depths <3 mm and became substantial (>1 mm), at 5 mm depth. Largest error (and non-linearity) was in inter-collicular regions, demonstrating where the Euclidean cylinder method is least appropriate.

### 3.2 – Depth profiles of visual stimulation- and saccade-evoked activity in SC

In superficial SC, our level-set depth was very similar to the original Euclidean depth. Our measurements of the BOLD response as a function of depth confirmed similar depth profiles, including the previous finding that the mean depth profiles across subjects for saccadic activity was shifted deeper into SC compared to those obtained from visual stimulation. This is consistent with the functional organization of SC delineated in animal models (Fuchs, Kaneko, & Scudder, 1985; Sparks & Hartwich-Young, 1989; Wurtz & Albano, 1980). The profiles generated using our new method did not show significant differences from those utilizing the cylinder method (Fig. 5); substantial differences in Euclidean and level-set depth metrics only emerge at depths >3 mm.

### 3.4 – Characterization of the PAG

Upon examination the gray matter probability maps of the PAG, we found that a large proportion of the dorsal midbrain is attributed a high probability value, even including superficial tissue that would be considered to be colliculus (Fig. 6A). However, after using a combination of the streamline depth kernels and Euclidean distance from CA, we were able to localize the PAG within this ambiguous region with a relatively small degree of uncertainty (Fig. 6B). The PAG for all subjects was generally able to be fitted with ellipses resulting in a root-mean-squared error of 0.65 mm (Fig. 6C), which did not show any significant variations within the slices of individual subjects. Ellipse parameters a, b, and ε (Fig. 6E) had coefficients-of-variation 0.37±0.08, 0.15±0.06, and 0.49±0.34, respectively, demonstrating that the elliptical fits were significantly more stable across subjects in the dorsal-ventral direction compared to the left-right direction. Rotation parameters were negligible. Visualization of the PAG contour and its associated surface, *S*_PAG_, reveals that this nucleus is not necessarily a uniform tube surrounding CA. Rather, its thickness, particularly along the left-right axis, varies along the length of CA (Fig. 6D and E).

The vertices of the PAG contour (*S*_PAG_) and our estimation of the contour boundary using ellipses (*S*_ellipse_) were very similar. Mean vertex separation, averaged across subjects, was 0.46±0.18 mm. A few vertices were widely separated; maximum separations were 7.2±5.0 mm. However, the fraction of these outlier vertices, with separations >2 mm was very small, 4.3±2.7%. Thus, the PAG is well characterized as a series of smooth ellipses that vary along the length of the CA. Our elliptical fitting procedure smoothed out many of the irregularities present in the actual PAG contours derived from the gray-matter probability maps (Fig. 6D), though some inhomogeneities are still observable in orthogonal cross-sections of *S*_ellipse_. However, particularly visible in a sagittal view, these “dimples” occur in the plane of the 2D ellipses, or in a direction that is normal to CA, reflecting some natural variance in the elliptical parameters along the length of CA. We would expect them to be alleviated by further refinement. Overall, our results confirmed the utility of Euclidean distance from CA as a reference depth metric for this region of the midbrain.

### 3.5 – Measurements of collicular tissue thickness

Thickness maps calculated using level-set physical distance from *S*_1_ to *S*_PAG_ showed that the tissue of the colliculi tends to be thickest at its crown, as expected, but also extending laterally, particularly the superior lateral edges for SC. The tissue in between left and right colliculi, the inter-collicular sulcus, was thinnest (Fig. 7A). Mean data across subjects for tissue thickness at the four peaks of SC and IC as well as the two sulci between left and right SC and IC are shown in Table 1. Interestingly, the variability of SC thickness was 2.77 mm of level-set depth, while IC thickness was significantly more stable at 0.97 mm. A similar pattern was observed for superior and inferior inter-collicular sulci, which had variabilities of 0.88 and 0.46 mm, respectively. For absolute thickness measures, there were no significant left-right thickness asymmetries. Thickness values at the inter-collicular sulci were very similar for inferior and superior colliculi. The normalized thickness metrics showed lower coefficients of variation in general. Left-right symmetry was also more clearly suggested by normalized thickness.

## 4 DISCUSSION

We compared an algebraic level-set scheme utilizing a normalized depth coordinate, *w*, to a Euclidean depth-mapping scheme for midbrain tissue between the surfaces of the brainstem and the cerebral aqueduct in humans. Previously, we used computational cylinders and Euclidean nearest-neighbor definitions of depth to estimate depth profiles of functional activity evoked by visual stimulation and attention (Katyal & Ress, 2014; Katyal et al., 2010; Savjani et al., 2018). However, the cylinder method was limited by contention of depth kernels, rendering it susceptible to oversampling deep tissue within individual colliculi and undersampling tissue in the inter-collicular sulcus. In contrast, the resulting non-linear coordinate system in native brain space using our level-set method is specifically adapted to the 3D shape of the tectum. Our novel streamline depth kernels never crossed paths by nature of the pseudopotential and compress or expand along their depth, which avoided mis-association between deep and superficial voxels using the cylinder method. The streamlines give a one-to-one association between the brainstem surface and the cerebral aqueduct (CA) at all spatial locations.

We also defined a new metric for physical tissue depth using streamline path distance. Both this level-set physical depth and the normalized *w* depth provide an analytic depth coordinate for each voxel. Additionally, the streamline depth kernels enable surface-based smoothing of the functional activity, provide mappings between the superficial surface *S*_1_) and deeper tissue, and allow quantitative data, such as gray matter probability, at different depths to be projected onto the surface (Dale, Fischl, & Sereno, 1999; Glasser et al., 2016). The parcellation of tissue into kernels provided depth relationships between deep and superficial tissue at a single spatial location, or over any functionally defined superficial surface. For example, the retinotopic organization of superficial superior colliculus (SC) (Cynader & Berman, 1972) could be related to the putative somatotopic organization in the deep layers of human SC suggested by experiments on animal models (Clemo & Stein, 1991; Jay & Sparks, 1987; Nagy, Kruse, Rottmann, Dannenberg, & Hoffmann, 2006; Wallace, Wilkinson, & Stein, 1996).

Overall, the streamline kernels are superior to the cylindrical kernels in their ability to create logical associations between tissue. However, we show that both the Euclidean and level-set depth metrics have their merits in appropriate regions of the midbrain. Usage of streamline depth kernels and level-set physical depth produced depth profiles of visual stimulation- and saccade-evoked activity in superficial SC similar to those previous obtained using cylindrical kernels and Euclidean distance from the surface of the brainstem, *d*_1_, as the reference depth metric (Fig. 5). Negligible differences between the methods in superficial tissue imply that either method works well here. On the other hand, the anatomical structure of the PAG is best defined using depth values according to Euclidean distance from CA, or *d*_2_ (Fig. 6). Thus, Euclidean distances from an appropriate reference surface are sufficient to delineate both superficial colliculus and the deep PAG.

Because of the emerging differences between the methods at greater depths within the midbrain, we predict that our streamlines and level-set depth metrics will provide the most utility in deeper tissue. Particularly in regard to the laminar structure of the colliculi, isosurfaces of our normalized *w* distance may provide a more accurate representation of the anatomical morphology of midbrain tissue compared to Euclidean definitions of tissue depth. Our normalized w depth may prove to be the most useful metric in regions of intermediate depth, namely deep SC and IC. In this case, confirmatory studies such as those involving reach-related activation (Linzenbold & Himmelbach, 2012; Liu, Duggan, Salt, & Cordeiro, 2011; Sparks & Hartwich-Young, 1989) are necessary.

We measured collicular thickness using our level-set definition of depth. While thickness measured in physical units (mm) varied between individuals, measurements in units of normalized depth demonstrated the least variability, testifying to the normalization that this metric provides between brains of different sizes. SC tended to have similar thickness, but greater variability compared to IC. While the mean thicknesses of left and right colliculi were not significantly different, left-right symmetry varied among individual subjects. Some subjects had differences in thickness greater than 2 mm, while others had differences <0.25 mm. The left-right differences in thickness were somewhat smaller for SC than IC.

However, from visual inspection of the PAG contour maps within the context of each subject’s anatomical volume, left-right asymmetry was not apparent. This disparity between expectation and qualitative observation, and our data is subject to two interpretations. First, what appears to be qualitatively symmetrical might not be in terms of path distance along nonlinear streamlines, which is not immediately evident purely from visual inspection. Second, and more importantly, the asymmetry tended to be an effect of which point on the BS surface was chosen to be the “peak” of each colliculus. Our method of choosing the point with the furthest straight-line distance to the base plane of each colliculus has an inherently Euclidean basis. Alternatively, taking the point with the greatest streamline distance to the base plane would have introduced a bias towards the very method we are attempting to test. It would also favor choosing points near the lateral edges where the streamlines are least perpendicular to the base plane. The desired output of the peak-finding algorithm is unclear at this point, especially when we have yet to fully characterize the location and lateral boundaries of the colliculi.

To the best of our knowledge, this work is the first to assess human collicular thicknesses *in vivo*, and these measurements, expanded to a larger pool of healthy brains, may be useful for gross delineation of neurodegenerative pathology that affect the colliculi such as glaucoma or aging (Crish, Sappington, Inman, Horner, & Calkins, 2010; Liu et al., 2011; Ouda & Syka, 2012). Though our data may appear to imply that some of our subjects suffered from unilateral degeneration, our method is one of many. Developing a way to accurately measure collicular tissue thickness depends on our ability to localize the entire boundary around these nuclei. Further comparison studies between high-resolution in vivo brain imaging and histology on postmortem brains, similar to (Loureiro et al., 2018), would be necessary to provide a complete analysis of collicular topology.

We strove for a method to relate observed function back to the anatomical structures from which they originate. Studying brain function within a structural context, as in a standardized atlas space (Evans et al., 1994; Mazziotta et al., 2001; Talairach & Tournoux, 1988), allows for localization of relevant structures and, consequently, generalizability across populations and studies. For example, the MNI atlas allows non-linear warping of 3D MRI brain scans from individuals onto a common template and has become a standard tool for reporting fMRI results in volume space. The PAG particularly lends itself for atlas comparisons, since this brain structure is relatively stable across subjects and with age (Keuken et al., 2017). Our experiments assumed that the tissue probability maps we used were well-suited to study the PAG. Usage of these maps has been validated using other deep brain nuclei including the thalamus, caudate, putamen, substantia nigra, subthalamic nucleus, red nucleus, and cerebellar dentate (Lorio et al., 2016). To our knowledge, this work is the first application of these methods to the PAG. Though these maps were not explicitly designed for localization of the PAG, small spatial uncertainty values (Fig. 6B) suggest that this application is appropriate. Our more rigorous quantitative method confirms the use of high gray-matter probability values for tissue classification. Furthermore, under the assumption that the PAG boundary can be fit by a continuous surface, our estimation scheme using ellipses worked well. We observed low root-mean-squared errors for individual elliptical fits, and low coefficients-of-variation for the ellipse parameters across subjects.

Further extension of our techniques to provide a fully three-dimensional coordinate system may utilize normalized depth and angle from a point of origin on the mid-sagittal plane through CA. This could potentially allow localization of midbrain nuclei such as the PAG using polar coordinate shapes such as dimpled or convex limaçons, which may provide a greater degree of accuracy than the simple ellipses we utilized in our work. Our efforts towards characterizing and subsequently reconstructing the PAG provide a first step in the direction towards creating a generalized atlas of midbrain nuclei (Fig. 6). Using CA as a central axis, as we have done, could form the basis of a quantitative atlas of the midbrain in a normalized coordinate space such as the MNI-305 (Mandal, Mahajan, & Dinov, 2012), but at higher spatial resolution. While we concentrated our evaluation of this depth-mapping scheme to the dorsal portion of midbrain, our method can be also effective for ventral midbrain, e.g., the red nucleus. It may be possible to interpolate the neuraxis between the exit of CA into the fourth ventricle and the geometric axis of the spinal cord at the base of the brainstem to facilitate similar methods in pons and medulla.

Our depth analysis assumed that the laminar structure of SC is well delineated by a depth metric derived from a combination of SC’s superficial surface and the CA. In order to minimize oversampling and overlapping depth projections, selecting an appropriate reference structure was key for defining depth in the midbrain. The CA runs rostro-caudally through the midbrain ventral to the colliculi. It emerges developmentally from the neural canal, the cavity in the center of the neural tube. The midbrain develops around the CA, thus positioning it as an important anatomical feature in the midbrain. Ideally, the resulting isosurfaces of *w* should be parallel to the laminae of the colliculi. However, our choice of surfaces to calculate depth coordinates was also a matter of convenience. Again, further comparison studies with histology on postmortem brains and high-resolution in vivo brain imaging (Loureiro et al., 2018) will be necessary to evaluate this assumption.

## 5 CONCLUSIONS

We established a series of morphologically reasonable methods for determining tissue depth in human SC and IC based solely on structural MRI. The level-set scheme provides coordinates for both normalized and physical depth that accommodate the complex anatomical structure of midbrain tissue, and the associated streamline method was able to distinguish function in the superficial layers of SC to the same accuracy as our previous Euclidean method. Euclidean distance from CA provides an accurate way to localize the PAG. We also believe that our work is the first analytic method for *in vivo* determination of anatomical tissue thickness in human midbrain. Because the streamline depth kernels logically follow the curvature of the tissue to establish unique relationships between all layers, our method should be useful to describe associations between tissue at different depths, such as between the deep multi-sensory layers of superior colliculus and the superficial retinotopically organized layers. Furthermore, depth-analysis studies using our normalized depth coordinate applied to a large population of subjects may enable a fully three-dimensional atlas of human brainstem.

## ACKNOWLEDGEMENTS

We gratefully acknowledge the support of lab members Elizabeth Halfen and Amanda Taylor in performing this work. This international collaborative work was supported by United States and German agencies: NIH grants R01 EB027586, K25 HL131997 and R01NS095933, BMBF CRCNS US-German Research Proposal Number 1822655.

## REFERENCES

Amidror, I. (2002). Scattered data interpolation methods for electronic imaging systems: a survey. Journal of Electronic Imaging, 11(2), 157. https://doi.org/10.1117/1.1455013

Ashburner, J., & Friston, K. J. (2005). Unified segmentation. NeuroImage, 26(3), 839–851. https://doi.org/10.1016/j.neuroimage.2005.02.018

Bajaj, C. L., Xu, G.-L., & Zhang, Q. (2008). Bio-molecule Surfaces Construction via a Higher-Order Level-Set Method. Journal of Computer Science and Technology, 23(6), 1026–1036. https://doi.org/10.1007/s11390-008-9184-1

Baumann, S., Griffiths, T. D., Rees, A., Hunter, D., Sun, L., & Thiele, A. (2010). Characterisation of the BOLD response time course at different levels of the auditory pathway in non-human primates. NeuroImage, 50(3), 1099–1108. https://doi.org/10.1016/j.neuroimage.2009.12.103

Baumann, S., Griffiths, T. D., Sun, L., Petkov, C. I., Thiele, A., & Rees, A. (2011). Orthogonal representation of sound dimensions in the primate midbrain. Nature Neuroscience, 14(4), 423–425. https://doi.org/10.1038/nn.2771

Cheung, M. M., Lau, C., Zhou, I. Y., Chan, K. C., Cheng, J. S., Zhang, J. W., … Wu, E. X. (2012). BOLD fMRI investigation of the rat auditory pathway and tonotopic organization. NeuroImage, 60(2), 1205–1211. https://doi.org/10.1016/j.neuroimage.2012.01.087

Clemo, H. R., & Stein, B. E. (1991). Receptive field properties of somatosensory neurons in the cat superior colliculus. The Journal of Comparative Neurology, 314(3), 534–544. https://doi.org/10.1002/cne.903140310

Crish, S. D., Sappington, R. M., Inman, D. M., Horner, P. J., & Calkins, D. J. (2010). Distal axonopathy with structural persistence in glaucomatous neurodegeneration. Proceedings of the National Academy of Sciences of the United States of America, 107(11), 5196–5201. https://doi.org/10.1073/pnas.0913141107

Cynader, M., & Berman, N. (1972). Receptive-field organization of monkey superior colliculus. Journal of Neurophysiology, 35(2), 187–201. https://doi.org/10.1152/jn.1972.35.2.187

Dale, A. M., Fischl, B., & Sereno, M. I. (1999). Cortical surface-based analysis: I. Segmentation and surface reconstruction. NeuroImage, 9(2), 179–194. https://doi.org/10.1006/nimg.1998.0395

De Martino, F., Moerel, M., van de Moortele, P.-F., Ugurbil, K., Goebel, R., Yacoub, E., & Formisano, E. (2013). Spatial organization of frequency preference and selectivity in the human inferior colliculus. Nature Communications, 4, 1386. https://doi.org/10.1038/ncomms2379

Eberly, D. (1999). Distance Between Point and Triangle in 3D. Retrieved from https://www.geometrictools.com/

Evans, A. C., Collins, D. L., Mills, S. R., Brown, E. D., Kelly, R. L., & Peters, T. M. (1994). 3D statistical neuroanatomical models from 305 MRI volumes. In IEEE Nuclear Science Symposium & Medical Imaging Conference (pp. 1813–1817). Publ by IEEE. https://doi.org/10.1109/nssmic.1993.373602

Feldon, P., & Kruger, L. (1970). Topography of the retinal projection upon the superior colliculus of the cat. Vision Research, 10(2), 135–143. Retrieved from http://www.ncbi.nlm.nih.gov/pubmed/5440778

Fuchs, A. F., Kaneko, C. R. S., & Scudder, C. A. (1985). Brainstem Control of Saccadic Eye Movements. Annual Review of Neuroscience, 8(1), 307–337. https://doi.org/10.1146/annurev.ne.08.030185.001515

Glasser, M. F., Coalson, T. S., Robinson, E. C., Hacker, C. D., Harwell, J., Yacoub, E., … Van Essen, D. C. (2016). A multi-modal parcellation of human cerebral cortex. Nature, 536(7615), 171–178. https://doi.org/10.1038/nature18933

Hagberg, G. E., Bause, J., Ethofer, T., Ehses, P., Dresler, T., Herbert, C., … Scheffler, K. (2017). Whole brain MP2RAGE-based mapping of the longitudinal relaxation time at 9.4T. NeuroImage, 144(Pt A), 203–216. https://doi.org/10.1016/j.neuroimage.2016.09.047

Iglesias, J. E., Van Leemput, K., Bhatt, P., Casillas, C., Dutt, S., Schuff, N., … Fischl, B. (2015). Bayesian segmentation of brainstem structures in MRI. NeuroImage, 113, 184–195. https://doi.org/10.1016/J.NEUROIMAGE.2015.02.065

Jay, M. F., & Sparks, D. L. (1987). Sensorimotor integration in the primate superior colliculus. I. Motor convergence. Journal of Neurophysiology, 57(1), 22–34. https://doi.org/10.1152/jn.1987.57.1.22

Jones, S. E., Buchbinder, B. R., & Aharon, I. (2000). Three-dimensional mapping of cortical thickness using Laplace’s equation. Human Brain Mapping, 11(1), 12–32. Retrieved from http://www.ncbi.nlm.nih.gov/pubmed/10997850

Katyal, S., & Ress, D. (2014). Endogenous attention signals evoked by threshold contrast detection in human superior colliculus. The Journal of Neuroscience : The Official Journal of the Society for Neuroscience, 34(3), 892–900. https://doi.org/10.1523/JNEUROSCI.3026-13.2014

Katyal, S., Zughni, S., Greene, C., & Ress, D. (2010). Topography of Covert Visual Attention in Human Superior Colliculus. Journal of Neurophysiology, 104(6), 3074–3083. https://doi.org/10.1152/jn.00283.2010

Keuken, M. C., Bazin, P. L., Backhouse, K., Beekhuizen, S., Himmer, L., Kandola, A., … Forstmann, B. U. (2017). Effects of aging on [Formula: see text], [Formula: see text], and QSM MRI values in the subcortex. Brain Struct Funct, 222(6) 2487–2505. https://doi.org/10.1007/s00429-016-1352-4

Khan, R., Zhang, Q., Darayan, S., Dhandapani, S., Katyal, S., Greene, C., … Ress, D. (2011). Surface-based analysis methods for high-resolution functional magnetic resonance imaging. Graphical Models, 73(6), 313–322. https://doi.org/10.1016/j.gmod.2010.11.002

Kim, J. H., Taylor, A., & Ress, D. (2017). Simple Signed-Distance Function Depth Calculation Applied to Measurement of the fMRI BOLD Hemodynamic Response Function in Human Visual Cortex (pp. 216–228). https://doi.org/10.1007/978-3-319-54609-4_16

Linzenbold, W., & Himmelbach, M. (2012). Signals from the Deep: Reach-Related Activity in the Human Superior Colliculus. Journal of Neuroscience, 32(40), 13881–13888. https://doi.org/10.1523/JNEUROSCI.0619-12.2012

Liu, M., Duggan, J., Salt, T. E., & Cordeiro, M. F. (2011). Dendritic changes in visual pathways in glaucoma and other neurodegenerative conditions. Experimental Eye Research, 92(4), 244–250. https://doi.org/10.1016/J.EXER.2011.01.014

Loftus, W. C., Malmierca, M. S., Bishop, D. C., & Oliver, D. L. (2008). The cytoarchitecture of the inferior colliculus revisited: a common organization of the lateral cortex in rat and cat. Neuroscience, 154(1), 196–205. https://doi.org/10.1016/j.neuroscience.2008.01.019

Lorio, S., Fresard, S., Adaszewski, S., Kherif, F., Chowdhury, R., Frackowiak, R. S., … Draganski, B. (2016). New tissue priors for improved automated classification of subcortical brain structures on MRI. NeuroImage, 130, 157–166. https://doi.org/10.1016/j.neuroimage.2016.01.062

Loureiro, J. R., Hagberg, G. E., Ethofer, T., Erb, M., Bause, J., Ehses, P., … Himmelbach, M. (2017). Depth-dependence of visual signals in the human superior colliculus at 9.4 T. Human Brain Mapping, 38(1), 574–587. https://doi.org/10.1002/hbm.23404

Loureiro, J. R., Himmelbach, M., Ethofer, T., Pohmann, R., Martin, P., Bause, J., … Hagberg, G. E. (2018). In-vivo quantitative structural imaging of the human midbrain and the superior colliculus at 9.4T. NeuroImage, 177, 117–128. https://doi.org/10.1016/J.NEUROIMAGE.2018.04.071

Malmierca, M. S., Izquierdo, M. A., Cristaudo, S., Hernandez, O., Perez-Gonzalez, D., Covey, E., & Oliver, D. L. (2008). A Discontinuous Tonotopic Organization in the Inferior Colliculus of the Rat. Journal of Neuroscience, 28(18), 4767–4776. https://doi.org/10.1523/JNEUROSCI.0238-08.2008

Mandal, P. K., Mahajan, R., & Dinov, I. D. (2012). Structural brain atlases: design, rationale, and applications in normal and pathological cohorts. Journal of Alzheimer’s Disease : JAD, 31 Suppl 3(0 3), S169–88. https://doi.org/10.3233/JAD-2012-120412

Marques, J. P., Kober, T., Krueger, G., van der Zwaag, W., Van de Moortele, P. F., & Gruetter, R. (2010). MP2RAGE, a self bias-field corrected sequence for improved segmentation and T1-mapping at high field. NeuroImage, 49(2), 1271–1281. https://doi.org/10.1016/j.neuroimage.2009.10.002

Mazziotta, J., Toga, A., Evans, A., Fox, P., Lancaster, J., Zilles, K., … Mazoyer, B. (2001, August 29). A probabilistic atlas and reference system for the human brain: International Consortium for Brain Mapping (ICBM). Philosophical Transactions of the Royal Society B: Biological Sciences. Royal Society. https://doi.org/10.1098/rstb.2001.0915

Meredith, M. A., & Stein, B. E. (1986). Visual, auditory, and somatosensory convergence on cells in superior colliculus results in multisensory integration. Journal of Neurophysiology, 56(3), 640–662. https://doi.org/10.1152/jn.1986.56.3.640

Moerel, M., De Martino, F., Uğurbil, K., Yacoub, E., & Formisano, E. (2015). Processing of frequency and location in human subcortical auditory structures. Scientific Reports, 5(1), 17048. https://doi.org/10.1038/srep17048

Nagy, A., Kruse, W., Rottmann, S., Dannenberg, S., & Hoffmann, K.-P. (2006). Somatosensory-Motor Neuronal Activity in the Superior Colliculus of the Primate. Neuron, 52(3), 525–534. https://doi.org/10.1016/J.NEURON.2006.08.010

Ouda, L., & Syka, J. (2012). Immunocytochemical profiles of inferior colliculus neurons in the rat and their changes with aging. Frontiers in Neural Circuits, 6, 68. https://doi.org/10.3389/fncir.2012.00068

Ress, D. B., Harlow, M. L., Marshall, R. M., & McMahan, U. J. (2004). Methods for generating high-resolution structural models from electron microscope tomography data. Structure. Cell Press. https://doi.org/10.1016/j.str.2004.07.022

Ress, D., & Chandrasekaran, B. (2013). Tonotopic organization in the depth of human inferior colliculus. Frontiers in Human Neuroscience, 7, 586. https://doi.org/10.3389/fnhum.2013.00586

Robinson, D. A. (1972). Eye movements evoked by collicular stimulation in the alert monkey. Vision Research, 12(11), 1795–1808. Retrieved from http://www.ncbi.nlm.nih.gov/pubmed/4627952

Savjani, R. R., Katyal, S., Halfen, E., Kim, J. H., & Ress, D. (2018). Polar-angle representation of saccadic eye movements in human superior colliculus. NeuroImage, 171, 199–208. https://doi.org/10.1016/J.NEUROIMAGE.2017.12.080

Schneider, K., & Kastner, S. (2010). Sustained spatial attention in the human lateral geniculate nucleus and superior colliculus. Journal of Vision, 7(9), 784–784. https://doi.org/10.1167/7.9.784

Schreiner, C. E., & Langner, G. (1997). Laminar fine structure of frequency organization in auditory midbrain. Nature, 388(6640), 383–386. https://doi.org/10.1038/41106

Shajan, G., Kozlov, M., Hoffmann, J., Turner, R., Scheffler, K., & Pohmann, R. (2014). A 16-channel dual-row transmit array in combination with a 31-element receive array for human brain imaging at 9.4 T. Magnetic Resonance in Medicine, 71(2), 870–879. https://doi.org/10.1002/mrm.24726

Singh, V., Pfeuffer, J., Zhao, T., & Ress, D. (2017). Evaluation of spiral acquisition variants for functional imaging of human superior colliculus at 3T field strength. Magnetic Resonance in Medicine, 79(4), 1931–1940. https://doi.org/10.1002/mrm.26845

Sparks, D. L., & Hartwich-Young, R. (1989). The deep layers of the superior colliculus. Reviews of Oculomotor Research, 3, 213–255. Retrieved from http://www.ncbi.nlm.nih.gov/pubmed/2486324

Sprague, J. M., & Meikle, T. H. (1965). THE ROLE OF THE SUPERIOR COLLICULUS IN VISUALLY GUIDED BEHAVIOR. Experimental Neurology, 11, 115–146. Retrieved from http://www.ncbi.nlm.nih.gov/pubmed/14272555

Talairach, J., & Tournoux, P. (1988). Co-planar Stereotaxic Atlas of the Human Brain. Stuttgart, Germany: Thieme Publishers.

Waehnert, M. D., Dinse, J., Weiss, M., Streicher, M. N., Waehnert, P., Geyer, S., … Bazin, P.-L. (2014). Anatomically motivated modeling of cortical laminae. NeuroImage, 93, 210–220. https://doi.org/10.1016/J.NEUROIMAGE.2013.03.078

Wallace, M. T., Wilkinson, L. K., & Stein, B. E. (1996). Representation and integration of multiple sensory inputs in primate superior colliculus. Journal of Neurophysiology, 76(2), 1246–1266. https://doi.org/10.1152/jn.1996.76.2.1246

Wurtz, R. H., & Albano, J. E. (1980). Visual-Motor Function of the Primate Superior Colliculus. Annual Review of Neuroscience, 3(1), 189–226. https://doi.org/10.1146/annurev.ne.03.030180.001201

Xu, G., Pan, Q., & Bajaj, C. L. (2006). Discrete surface modelling using partial differential equations. Computer Aided Geometric Design, 23(2), 125–145. https://doi.org/10.1016/J.CAGD.2005.05.004

Yarnykh, V. L. (2010). Optimal radiofrequency and gradient spoiling for improved accuracy of T1 and B1 measurements using fast steady-state techniques. Magnetic Resonance in Medicine, 63(6), 1610–1626. https://doi.org/10.1002/mrm.22394

Yarnykh, V., & Yuan, C. (2004). Actual flip angle imaging in the pulsed steady state. Proc Intl Soc Mag Reson Med, 11(1), 2004–2004. https://doi.org/10.1002/mrm.21120

Yushkevich, P. A., Piven, J., Hazlett, H. C., Smith, R. G., Ho, S., Gee, J. C., & Gerig, G. (2006). User-guided 3D active contour segmentation of anatomical structures: Significantly improved efficiency and reliability. NeuroImage, 31(3), 1116–1128. https://doi.org/10.1016/J.NEUROIMAGE.2006.01.015

